## Supplementary Material for "Patterns of recent natural selection on genetic loci associated with sexually differentiated human body size and shape phenotypes"

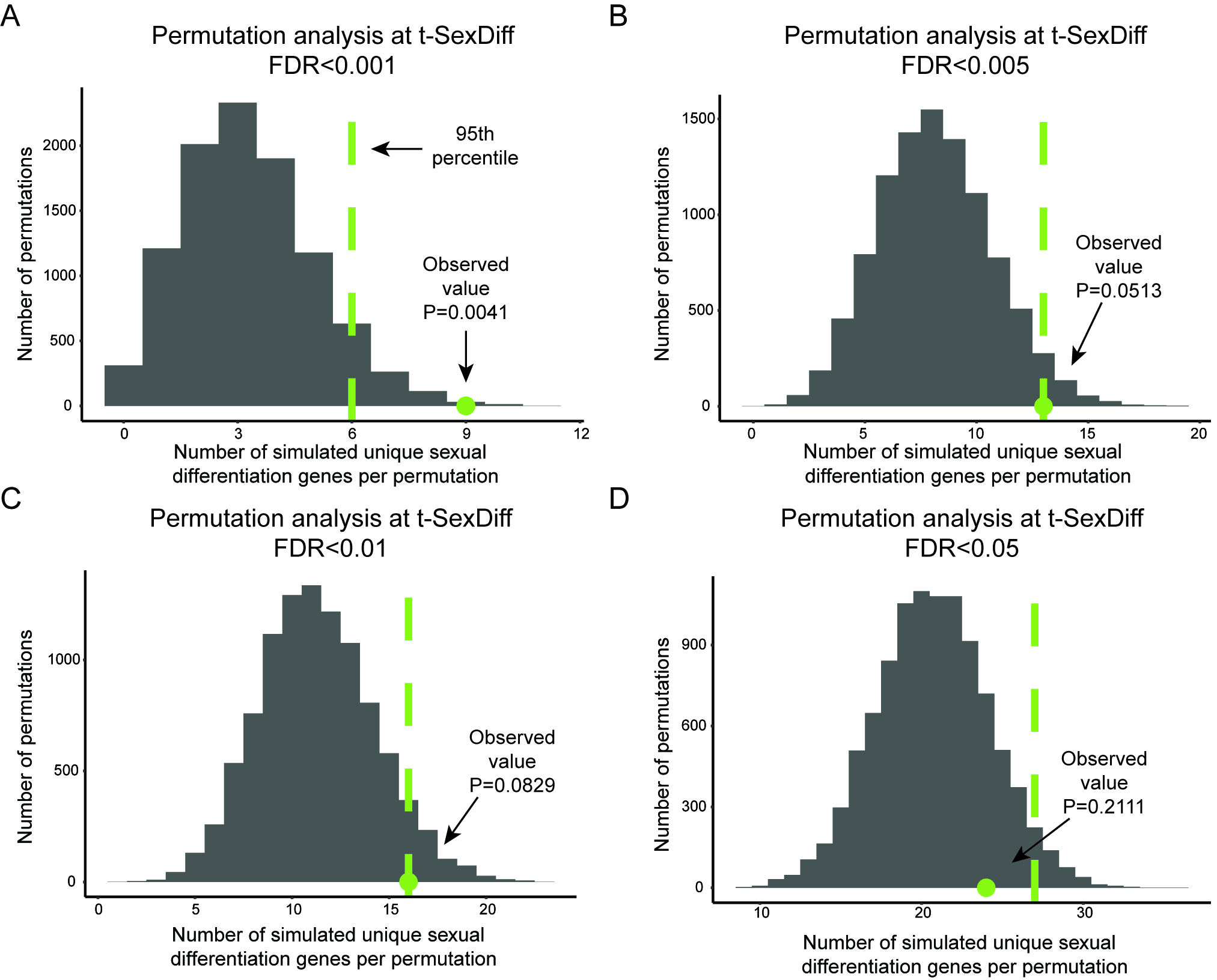

**S1 Fig. Permutation enrichment distribution at each FDR threshold.** Permutation analysis of the number of genes involved in sexual differentiation for all anthropometric SNPs at every FDR threshold. Data are the frequency of distribution of our results for 10,000 permuted data sets. The empirical P-value represents the probability that the observed value of sexual differentiation genes from our SexDiff-associated SNP pool is equal to or greater than those from a randomly selected set.

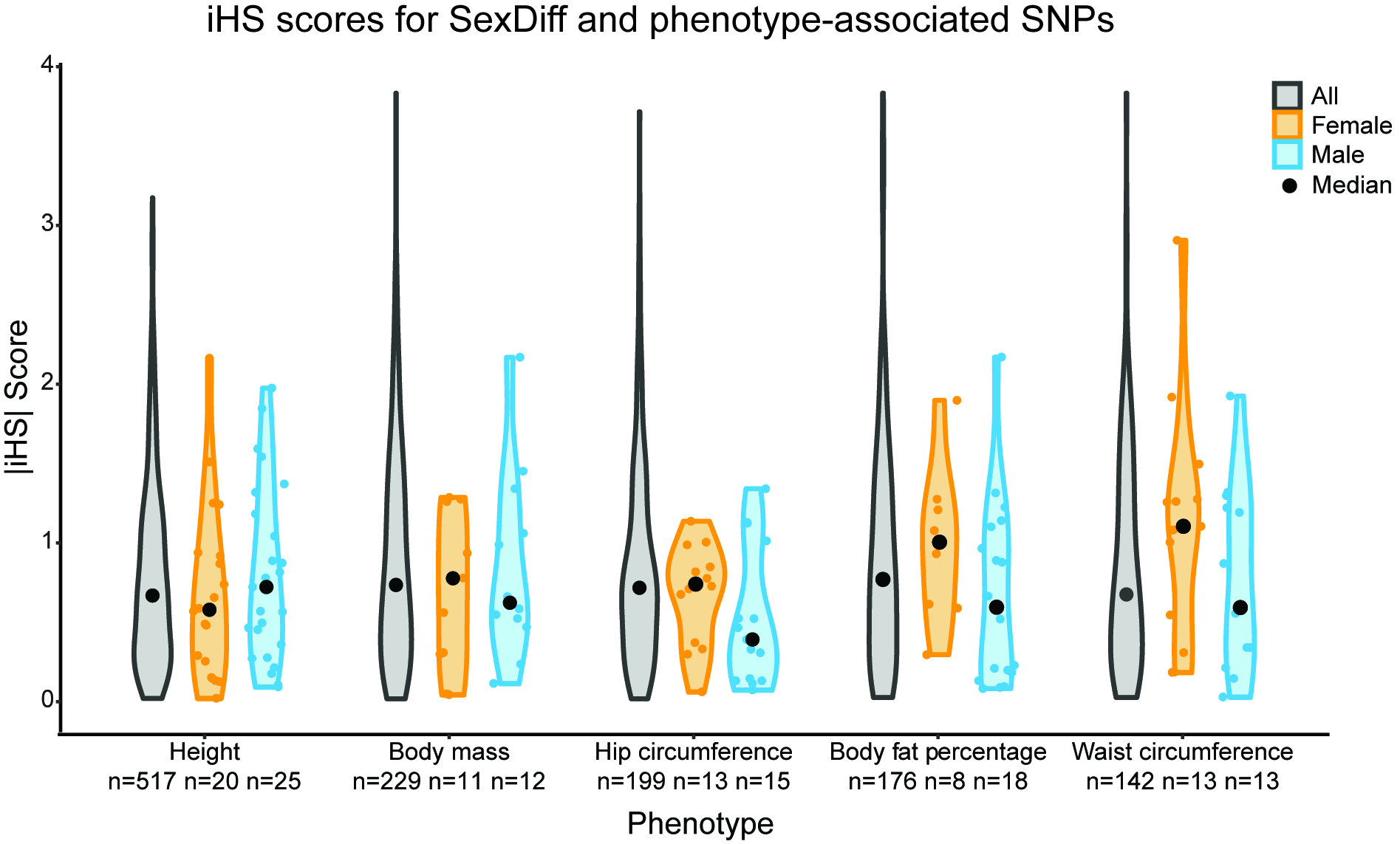

**S2 Fig. Sex-specific iHS scores for anthropometric SexDiff and phenotype-associated SNPs.** The iHS distributions for each set of female and male SexDiff pruned SNPs were compared to those for the corresponding phenotype-association set using a permutation analysis. None of the distributions were significantly different.

**S1 Table:** Observed number of SexDiff-associated SNPs at each FDR threshold for every phenotype.

| Phenotype | Number of phenotype-associated SNPs | Number of SexDiff-associated SNPs | | | | Ratio^a^ |
| --- | --- | --- | --- | --- | --- | --- |
|  |  | FDR  0.05 | FDR  0.01 | FDR  0.005 | FDR  0.001 |  |
| Height | 67738 | 9654 | 4242 | 3803 | 677 | 0.0010 |
| Body mass | 15669 | 3589 | 1941 | 1102 | 541 | 0.0345 |
| Hip circumference | 12580 | 3822 | 1991 | 1422 | 808 | 0.0642 |
| Body fat percentage | 10538 | 3488 | 1727 | 1034 | 439 | 0.0417 |
| Waist circumference | 7675 | 2479 | 897 | 742 | 551 | 0.0718 |

^a^Ratio of SexDiff-associated SNPs at the FDR threshold of 0.001 to the number of phenotype-associated SNPs

**S2 Table:** Observed unique sexual differentiation genes (SDG) and total number of genes for SexDiff-associated SNPs and Non SexDiff-associated SNPs

| FDR threshold | Number of unique sexual differentiation genes | | | | | | Ratio^b^ |
| --- | --- | --- | --- | --- | --- | --- | --- |
|  | SexDiff-associated SNPs | | | Non SexDiff-associated SNPs | | |  |
|  | SDG | Total Genes | Proportion^a^ | SDG | Total Genes | Proportion^a^ |  |
| 0.001 | 9 | 162 | 0.0555 | 52 | 2544 | 0.0204 | 2.7206 |
| 0.005 | 13 | 396 | 0.0328 | 51 | 2499 | 0.0204 | 1.6078 |
| 0.01 | 16 | 545 | 0.0294 | 49 | 2437 | 0.0201 | 1.4627 |
| 0.05 | 24 | 1005 | 0.0239 | 47 | 2269 | 0.0207 | 1.1546 |

^a^Proportion of SDG genes observed in each SNP group ^b^Ratio of proportion of SDG genes observed in the group of SexDiff-associated SNPs to the proportion of SDG genes observed in the group of non SexDiff-associated SNPs

**S3 Table:** Observed log_2_ ratio of female to male beta values and p-values for each set of Female SexDiff-associated SNPs

| Phenotype | #SNPs^a^ | Mean  log_2_(ratio)^b^ | P-value to zero^c^ | FDR to zero | P-value to phenotype-associated SNPs^d^ | FDR to phenotype-associated SNPs |
| --- | --- | --- | --- | --- | --- | --- |
| Height | 21 | 0.2403 | 4.8x10^-10^ | 4.8x10^-9^ | <0.001 | 0.001 |
| Body mass | 11 | 0.2331 | 9.8x10^-7^ | 1.6x10^-6^ | <0.001 | 0.001 |
| Hip circumference | 13 | 0.1739 | 1.9x10^-6^ | 2.7x10^-6^ | <0.001 | 0.001 |
| Body fat percentage | 9 | 0.8272 | 3.3x10^-5^ | 3.3x10^-5^ | <0.001 | 0.001 |
| Waist circumference | 14 | 0.0222 | 5.1x10^-6^ | 6.4x10^-6^ | <0.001 | 0.001 |

^a^Number of pruned SexDiff-associated SNPs at an FDR threshold of 0.001 ^b^Mean log_2_ ratio of female trait effect size to the male trait effect size ^c^One-sided t-test P-value comparing distribution of the log_2_(ratio) ^d^Permutation P-value of the probability that the mean log_2_(ratio) could be observed by chance when compared to phenotype-associated SNPs

**S4 Table:** Observed log_2_ ratio of female to male beta values and p-values for each set of Male SexDiff-associated SNPs

| Phenotype | #SNPs^a^ | Mean  log_2_(ratio)^b^ | P-value to zero^c^ | FDR to zero | P-value to phenotype-associated SNPs^d^ | FDR to phenotype-associated SNPs |
| --- | --- | --- | --- | --- | --- | --- |
| Height | 25 | 0.2403 | 2.3x10^-7^ | 5.8x10^-7^ | <0.001 | 0.001 |
| Body mass | 12 | 0.2331 | 2.2x10^-7^ | 5.8x10^-7^ | <0.001 | 0.001 |
| Hip circumference | 15 | 0.1739 | 7.8x10^-8^ | 3.9x10^-7^ | <0.001 | 0.001 |
| Body fat percentage | 18 | 0.8272 | 8.0x10^-7^ | 1.6x10^-6^ | <0.001 | 0.001 |
| Waist circumference | 16 | 0.0222 | 1.8x10^-5^ | 2.0x10^-5^ | <0.001 | 0.001 |

^a^Number of pruned SexDiff-associated SNPs at an FDR threshold of 0.001 ^b^Mean log_2_ ratio of female trait effect size to the male trait effect size ^c^One-sided t-test P-value comparing distribution of the log_2_(ratio) ^d^Permutation P-value of the probability that the mean log_2_(ratio) could be observed by chance when compared to phenotype-associated SNPs

**S5 Table:** Observed trait-SDS and permutation P-values for each set of Female SexDiff-associated SNPs and Male SexDiff-associated SNPs permuted against phenotype-associated SNPs.

| Phenotype | Female | | | | Male | | | |
| --- | --- | --- | --- | --- | --- | --- | --- | --- |
|  | #SNPs^a^ | trait-SDS^b^ | P-value to phenotype-associated SNPs^c^ | FDR | #SNPs^a^ | trait-SDS^b^ | P-value to phenotype-associated SNPs^c^ | FDR |
| Height | 21 | 0.2403 | 0.6608 | 0.8246 | 25 | 0.0540 | 0.6944 | 0.8246 |
| Body mass | 11 | 0.2331 | 0.7864 | 0.8246 | 12 | 0.2359 | 0.7408 | 0.8246 |
| Hip circumference | 13 | 0.1739 | 0.4688 | 0.8246 | 15 | 0.7068 | 0.1208 | 0.6040 |
| Body fat percentage | 9 | **0.8272** | **0.0038** | **0.0380** | 18 | 0.1146 | 0.3046 | 0.8246 |
| Waist circumference | 14 | 0.0222 | 0.8246 | 0.8246 | 16 | 0.2161 | 0.6040 | 0.8246 |

^a^Number of pruned SexDiff-associated SNPs at an FDR threshold of 0.001 ^b^Average trait-SDS score of pruned set of SexDiff-associated SNPs ^c^Permutation P-value of the probability that the trait-SDS score for each sex could be observed by chance when compared to phenotype-associated SNPs

**S6 Table:** Observed trait-SDS and permutation P-values for each set of Female SexDiff-associated SNPs and Male SexDiff-associated SNPs permuted against phenotype-associated SNPs matched for minor allele frequency.

| Phenotype | Female | | | | Male | | | |
| --- | --- | --- | --- | --- | --- | --- | --- | --- |
|  | #SNPs^a^ | trait-SDS^b^ | P-value to phenotype-associated SNPs^c^ | FDR | #SNPs^a^ | trait-SDS^b^ | P-value to phenotype-associated SNPs^c^ | FDR |
| Height | 21 | 0.2403 | 0.7048 | 0.7987 | 25 | 0.0540 | 0.6362 | 0.7987 |
| Body mass | 11 | 0.2331 | 0.6390 | 0.7987 | 12 | 0.2359 | 0.6278 | 0.7987 |
| Hip circumference | 13 | 0.1739 | 0.7188 | 0.7987 | 15 | 0.7068 | 0.0576 | 0.2880 |
| Body fat percentage | 9 | **0.8272** | **0.0028** | **0.0280** | 18 | 0.1146 | 0.3674 | 0.7987 |
| Waist circumference | 14 | 0.0222 | 0.6616 | 0.7987 | 16 | 0.2161 | 0.8414 | 0.8414 |

^a^Number of pruned SexDiff-associated SNPs at an FDR threshold of 0.001 ^b^Average trait-SDS score of pruned set of SexDiff-associated SNPs ^c^Permutation P-value of the probability that the trait-SDS score for each sex could be observed by chance when compared to phenotype-associated SNPs matched for minor allele frequency

**S7 Table:** Observed average |iHS| scores and permutation P-values for each set of Female SexDiff-associated SNPs and Male SexDiff-associated SNPs permuted against phenotype-associated SNPs.

| Phenotype | Female | | | | Male | | | |
| --- | --- | --- | --- | --- | --- | --- | --- | --- |
|  | #SNPs^a^ | \|iHS\| | P-value to phenotype-associated SNPs^c^ | FDR | #SNPs^a^ | \|iHS\| | P-value to phenotype-associated SNPs^c^ | FDR |
| Height | 20 | 0.6586 | 0.7989 | 0.989 | 25 | 0.7822 | 0.4291 | 0.989 |
| Body mass | 11 | 0.6705 | 0.8368 | 0.989 | 12 | 0.8284 | 0.5659 | 0.989 |
| Hip circumference | 13 | 0.6547 | 0.8333 | 0.989 | 15 | 0.4975 | 0.9890 | 0.989 |
| Body fat percentage | 8 | 0.9729 | 0.3266 | 0.989 | 18 | 0.6570 | 0.9321 | 0.989 |
| Waist circumference | 13 | 1.1015 | 0.0770 | 0.770 | 13 | 0.7548 | 0.6384 | 0.989 |

^a^Number of pruned SexDiff-associated SNPs at an FDR threshold of 0.001 ^b^Average |iHS| score of pruned set of SexDiff-associated SNPs ^c^Permutation P-value of the probability that the |iHS| score for each sex could be observed by chance when compared to phenotype-associated SNPs

| Phenotype | Female | | | | Male | | | |
| --- | --- | --- | --- | --- | --- | --- | --- | --- |
|  | #SNPs^a^ | #Inter-genic SNPs | P-value to phenotype-associated SNPs^b^ | FDR | #SNPs^a^ | #Inter-genic SNPs | P-value to phenotype-associated SNPs^b^ | FDR |
| Height | 21 | 9 | 0.1498 | 0.2497 | 25 | 11 | 0.0998 | 0.2497 |
| Body mass | 11 | 5 | 0.1168 | 0.2497 | 12 | 4 | 0.4978 | 0.5531 |
| Hip circumference | 13 | 4 | 0.6204 | 0.6204 | 15 | 8 | 0.0178 | 0.0890 |
| Body fat percentage | 9 | 4 | 0.2312 | 0.2890 | 18 | 8 | 0.2186 | 0.2890 |
| Waist circumference | 14 | 6 | 0.1470 | 0.2497 | 16 | 8 | 0.0048 | 0.0480 |

**S8 Table:** Observed number of intergenic SNPs and permutation P-values for each set of Female SexDiff-associated SNPs and Male SexDiff-associated SNPs permuted against phenotype-associated SNPs.

^a^Number of pruned SexDiff-associated SNPs at an FDR threshold of 0.001 ^b^Permutation P-value of the probability that the number of intergenic SNPs in each set of SexDiff-associated SNPs could be observed by chance when compared to phenotype-associated SNPs

**S9 Table:** Phenotype information

| Phenotype | Phenotype description^a^ | Code^a^ | Sample sizes | | rho^b^ |
| --- | --- | --- | --- | --- | --- |
|  |  |  | # Females | #Males |  |
| Height | standing height | 50_irnt | 193,785 | 166,603 | 0.44617 |
| Body mass | weight | 21002_irnt | 193,627 | 166,489 | 0.37463 |
| Hip circumference | hip circumference | 49_irnt | 193,814 | 166,707 | 0.33003 |
| Body fat percentage | body fat percentage | 23099_irnt | 190,991 | 163,637 | 0.34575 |
| Waist circumference | waist circumference | 48_irnt | 193,828 | 166,736 | 0.31999 |

^a^As referred to in the Neale lab manifest released on July 31, 2018 ^b^ Correlation for each phenotype calculated as the Spearman rank correlation coefficient between beta values of men and women

**S10 Table:** Observed trait-SDS for each set of pruned phenotype-associated SNP groups.

| Phenotype | # Pruned SNPs | tSDS |
| --- | --- | --- |
| Height | 532 | 0.1325 |
| Body mass | 239 | 0.1496 |
| Hip circumference | 210 | 0.2123 |
| Body fat percentage | 181 | -0.1244 |
| Waist circumference | 147 | 0.0784 |
